# Equilibrium Tension and Compression Mechanical Properties of the Human Uterus

**DOI:** 10.1101/2024.04.25.591208

**Authors:** Shuyang Fang, Camilo A. Duarte-Cordon, Daniella M. Fodera, Lei Shi, Xiaowei Chen, Arnold Advincula, Joy Vink, Christine Hendon, Kristin M. Myers

## Abstract

A successful pregnancy relies on the proper cellular, biochemical, and mechanical functions of the uterus. A comprehensive understanding of uterine mechanical properties during pregnancy is key to understanding different gynecological and obstetric disorders such as preterm birth, placenta accreta, leiomyoma, and endometriosis. This study sought to characterize the macro-scale equilibrium material behaviors of the human uterus in non-pregnancy and late pregnancy under both compressive and tensile loading. Fifty human uterine specimens from 16 patients (8 nonpregnant [NP] and 8 pregnant [PG]) were tested using spherical indentation and uniaxial tension coupled with digital image correlation (DIC). A three-level incremental load–hold protocol was applied to both tests. A microstructurally-inspired material model considering fiber architecture was applied to this dataset. Inverse finite element analysis (IFEA) was then performed to generate a single set of mechanical parameters to describe compressive and tensile behaviors. The freeze-thaw effect on uterine macro mechanical properties was also evaluated. PG tissue exhibits decreased overall stiffness and increased fiber network extensibility compared to NP uterine tissue. Under indentation, ground substance compressibility was similar between NP and PG uterine tissue. In tension, the fiber network of the PG uterus was found to be more extensible and dispersed than in nonpregnancy. Lastly, a single freeze-thaw cycle did not systematically alter the macro-scale material behavior of the human uterus.

## 1. Introduction

The uterus is a critical organ in human pregnancy. In nonpregnancy, the uterus is a pear-shaped thick-walled muscular organ. In an uncomplicated pregnancy, the uterus undergoes significant growth and remodeling, expanding and stretching to more than 500 times its original volume-carrying capacity to accommodate the growing fetus and amniotic sac.[1, 2] At term, initiated by a combination of biochemical and biomechanical signals, the uterus must contract rapidly and intensely to enable safe delivery of the baby. Premature activation of uterine contraction is one known cause of preterm labor and birth (PTB).[3] PTB remains the leading cause of neonatal death, with one in every ten babies born preterm yearly around the globe.[4, 5, 6] Conversely, late activation of uterine contraction contributes to long gestational lengths greater than 40 weeks and prolonged labor, which often require pharmocologic (e.g., pitocin) or surgical (e.g. Cesarean section) interventions[7] Increased uterine stretch, resulting in uterine overdistension, is thought to be associated with the premature onset of contractions and higher rates of PTB among women carrying twins or excessive amniotic fluid[8, 9] Therefore, timely uterine remodeling and appropriate biomechanical properties are critical for ensuring a healthy pregnancy and labor. Characterizing the normal and abnormal remodeling of the uterus from nonpregnant (NP) to pregnant (PG) states can contribute to an increased understanding of pre-term and post-term labor mechanisms.

Previously, the mechanical properties of the human uterus have been evaluated *in vivo* with diffusion MRI and shear wave elastography and *ex vivo* with traditional mechanical tests under compression, indentation, and tension [10, 11, 12, 13, 14, 15] The mechanical properties of the human uterus are nonlinear, anisotropic, and tension-compression asymmetric. This complexity reflects the uterus’s unique structure, composed of architected smooth muscle cell fascicles sheathed in the collagen-rich extracellular matrix.[16, 17, 18] Previous *ex vivo* mechanical testing studies of the uterus reported tensile engineering stress–strain relationships between PG and NP uteri, and found a less steep slope for PG tissue.[10] A previous study by our lab used a fibrous material model to describe the equilibrium compressive properties of the uterus under indentation and found the uterine tissue anisotropic.[12]

Building on previous studies, this work seeks to characterize the tensile and compressive properties of the human uterus under large strains, as seen in gestation. In particular, this study focuses on the thick, muscular middle myometrial layer of the uterus, as it bears the greatest mechanical loads[17]. A single set of material parameters from a previously described microstructurally-inspired, hyperelastic material constitutive model describes the tensile and compressive uterine properties.[19] Fifty uterine specimens from 16 patients (8 nonpregnant and 8 pregnant) were studied. To obtain material properties, fiber structural data from previous Optical Coherence Tomography (OCT) imaging studies were integrated into the model a priori,[12, 16] and spherical indentation and uniaxial tension tests were performed to measure tissue material response under different loading conditions. Then, inverse finite element analysis was performed to determine the material parameters for each specimen.[12, 16] While the time-dependent properties of biological tissues play an important role in their material functions, this study first focuses on the equilibrium properties. Future modeling studies will use this equilibrium modeling foundation to characterize uterine material behavior fully.

## 2. Methods

A workflow for mechanical testing and data analysis was previously developed to investigate the equilibrium material properties of the nonhuman primate (NHP) cervix.[20] In this section, the methods adopted from previous work are summarized, and those that differ are described in greater detail. Briefly, fifty human uterine specimens were mechanically tested under spherical indentation and uniaxial tension, coupled with digital image correlation (DIC). A fibrous anisotropic material model informed by imaging results was then applied to capture uterine material behavior. Lastly, an inverse finite element analysis was performed and the material model was validated to determine the uterine material properties.

### 2.1. Specimen Collection and Processing

Human uterine specimens were collected from 16 patients (*<* 50 years old) away from visible sites of pathology. Nonpregnant individuals underwent a total hysterectomy for non-cancerous indications, while pregnant patients underwent a cesarean hysterectomy in the early third trimester due to suspected abnormal placentation. Patient information, including age, race & ethnicity, surgical indication, and obstetric history, is listed in Table 1. This study was approved by the Institutional Review Board at Columbia University Irving Medical Center (CUIMC), and each subject provided written informed consent.

**Table 1:** Summary of Patient Characteristics.

| Patient ID | Age (yrs) | Race | Eth | G/P | Estimated Menstrual Cycle Stage | Gestational Age (wks+days) | Diagnosis & Pathological Findings | Prior Obstetric & Gynecologic History |
| --- | --- | --- | --- | --- | --- | --- | --- | --- |
| NP1 | 40 | AA | Not Hisp | 1/0 | Secretory | N/A | Leiomyoma, Poly-poid endometrium | D&C×1 |
| NP2 | 49 | Unk | Hisp | 0/0 | Proliferative | N/A | Leiomyoma, Chronic endometritis | None |
| NP3 | 44 | AA | Not Hisp | 2/2 | Late Secretory | N/A | Leiomyoma, Adenomyosis | C/S×1, VD×1 |
| NP4 | 47 | Unk | Unk | 2/2 | Weakly Proliferative – Perimenopausal | N/A | Leiomyoma, Adenomyosis, Chronic endometritis | VD×2 |
| NP5 | 41 | White | Not Hisp | 3/2 | None – Postmenopausal | N/A | Leiomyoma, Adenomyosis, Atrophic endometrium | VD×2, SAB×1, Current progesterone use |
| NP6 | 36 | Unk | Hisp | 3/3 | Proliferative | N/A | Leiomyoma, Adenomyosis, Prolapse (uterus & vagina) | VD×3 |
| NP7 | 43 | Unk | Hisp | 4/3 | Unknown | N/A | Adenomyosis, Endometriosis, Chronic endometritis | VD×3, SAB×1, Tubal ligation |
| NP8 | 48 | White | Not Hisp | 4/3 | Early Secretory | N/A | Leiomyoma, Adenomyosis, Serosal adhesions | SAB×1, C/S×2, VD×1, Placenta previa×1, Current progestin use |
| PG1 | 29 | AA | Not Hisp | 5/2 | N/A | 35 + 1 | Placenta percreta | C/S×1, D&C×1, SAB×1, EP×1 |
| PG2 | 39 | Asian | Not Hisp | 4/3 | N/A | 35 + 1 | Placenta increta, Placenta previa, Adenomyosis | C/S×2 |
| PG3 | 37 | White | Not Hisp | 3/2 | N/A | 34 | Placenta increta | D&C×1, VD×2 |
| PG4 | 39 | AA | Not Hisp | 11/2 | N/A | 34 | Placenta accreta | C/S×2 |
| PG5 | 33 | Unk | Hisp | 5/4 | N/A | 36 + 3 | Placenta accreta | C/S×2, SAB×1 |
| PG6 | 31 | Unk | Unk | 4/3 | N/A | 32 | Placenta percreta, placenta previa | C/S×3 |
| PG7 | 46 | White | Not Hisp | 8/6 | N/A | 35 + 5 | Placenta accreta | C/S×1, D&C×1, VBAC×2 |
| PG8 | 38 | Unk | Unk | 14/12 | N/A | 36 + 1 | Placenta increta | C/S×1, VBAC×6 |

[KEY] G ≡ Gravidity; P ≡ Parity; Eth ≡ Ethnicity; AA ≡ African-American; Unk ≡ Unknown; Hisp ≡ Hispanic; D&C ≡ dilation and curettage; C/S ≡ Cesarean section; VD ≡ Vaginal delivery; SAB ≡ spontaneous abortion; EP ≡ ectopic pregnancy; VBAC ≡ vaginal birth after cesarean

Our previous work on characterizing human uterine tissue anisotropic properties using spherical indentation [12] described the tissue collection and processing procedure performed for this study. In summary, each specimen slice measured 15-25 mm in both length and width and approximately 4 mm in thickness and was stored at -80℃. Prior to testing, frozen tissue was equilibrated at 4℃ overnight using phosphate-buffered saline (PBS) solution supplemented with 2 mM ethylenediaminetetraacetic acid (EDTA) to minimize tissue degradation. In our previous work, 14 specimens of a single NP patient and 13 specimens of a single PG patient were imaged using OCT and mechanically tested via spherical indentation. These 27 specimens were collected from three anatomical locations, anterior, fundus, and posterior, throughout the full uterine wall thickness (including endometrium, myometrium, and perimetrium). Because the mechanical properties of uterine tissue at the fundus were between the properties of the anterior and posterior tissue samples, all tissue tested in this study was taken from the fundus region. Additionally, because the thick center myometrial layer takes the primary mechanical role of the uterus during gestation, this study focuses on the myometrium. Hence, two fundus myometrium specimens per patient from 14 patients (7 NP and 7 PG) were added to the existing 22 myometrial specimens from Fang et al. [12]. Therefore, 50 specimens were studied in this work, including 24 NP samples from 8 patients and 26 PG samples from 8 patients.

### 2.2. Mechanical Testing

Each tissue specimen was mechanically tested in PBS+EDTA solution under indentation (6-mm diameter) followed by tension (Fig. 2(a-b)). The mechanical tests were performed using a universal testing machine (Instron, Inc., Norwood, MA) with a 5 N load cell (Instron, Inc., Norwood, MA, accuracy of 0.005 N). For indentation, a three-level load–hold protocol was performed with indentation depths prescribed as displacement—15%, 30%, 45% of the specimen thickness (Fig. 2(c)). After each ramp, the indenter was held in place for 480, 600, and 720 seconds, respectively, for the specimen to approach equilibrium. After indentation, each sample was allowed to recover and was processed for testing in tension. Specifically, the specimen was cut into rectangular strips with an approximate length-to-width ratio of 2. The length direction was determined by aligning the fiber direction (observed by OCT) with the tension loading direction. For each tension test, to prevent slippage of the sample during testing, the specimen was placed between metal tensile grips fastened with a section of water-resistant sandpaper (McMaster-Carr, Aurora, OH) using Krazy glue (KrazyGlue, High Point, NC). A three-level load–hold–unload protocol was performed with extensions prescribed as engineering strains—15%, 30%, 45% calculated as the grip-to-grip (G2G) distance divided by the specimen gage length (Fig. 2(c)). The engineering strain rate was set as 0.1%*·*second*^−^*^1^. At each hold, the specimen was kept in place for 30, 45, and 60 minutes to approach equilibrium. Before each test, the tissue was pre-tensioned to 1 mN to remove any slack. All specimens were speckled with water-insoluble ink (Chartpak, Inc., Leeds, MA) using an airbrush (Harder & Steenbeck GmbH & Co., Germany) and a Sprint Jet air compressor (Iwata Medea Inc., Portland, OR) to enable tracking of full-field displacements. One front camera and two orthogonal cameras (Point Gray Grasshopper, GRAS-50S5M-C75 mm, f/4 lens) were used for the indentation and tension tests, respectively, to track deformations (Fig.2(a-b)). Calibration images were taken for each test using a ruler with 1/16-inch gradations in the field of view (FOV). For both mechanical tests, force-displacement-time (N-mm-s) data were collected using the material tester software (Instron, Inc., Blue Hill version 3.11.1209).

**Figure 1:**
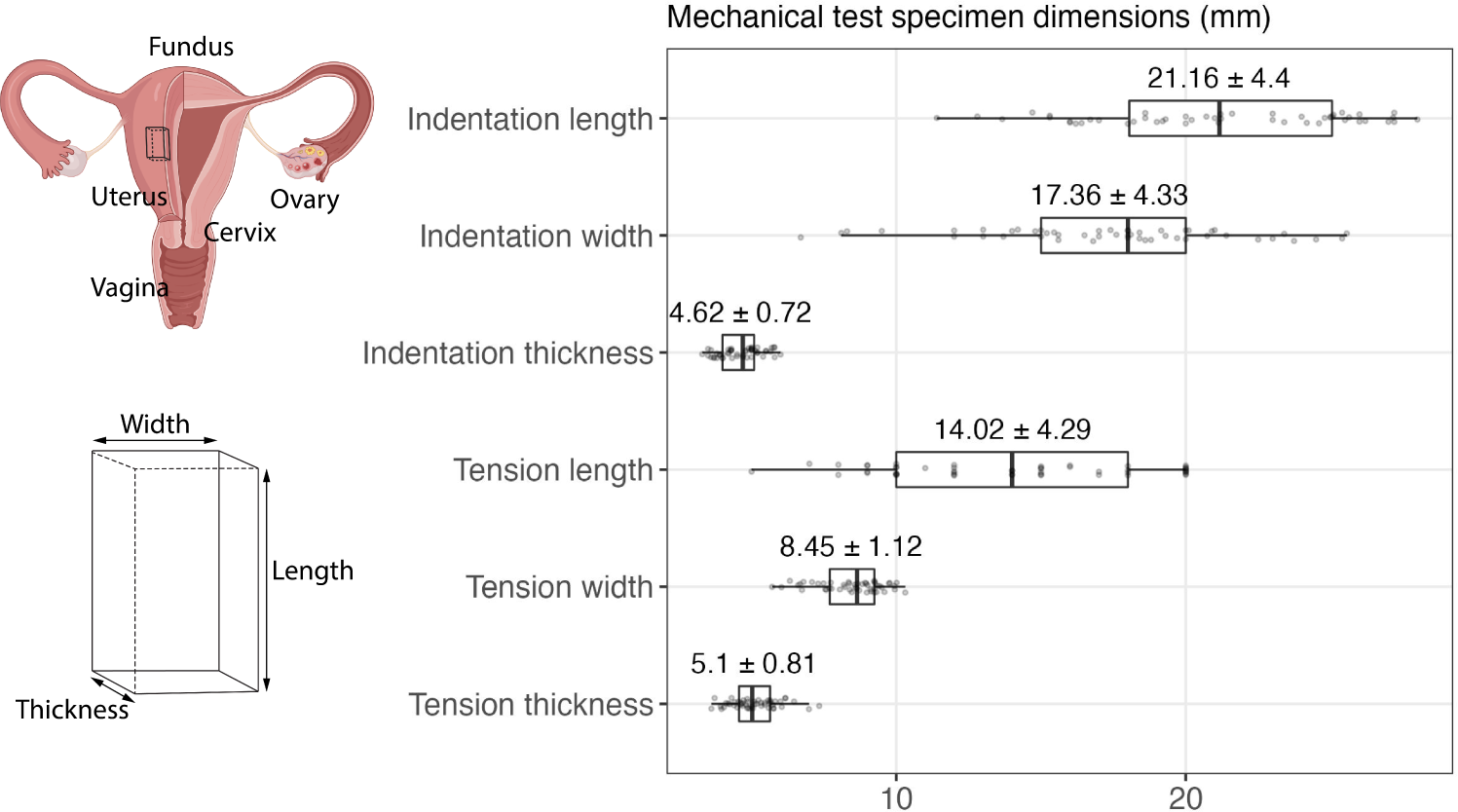
Specimen dimensions for mechanical tests. The specimens’ length, width, and thickness are plotted for indentation and tension tests. The mean *±* standard deviation values are calculated for all 50 specimens and labeled above each category’s box. During tension testing, one end of the specimen was affixed to the top tensile jaw, and the specimen was lowered into the bottom jaw. Therefore, the tensile specimen’s grip-to-grip (G2G) length was chosen and measured as integer numbers. The opaque black dots represent all data points.

**Figure 2:**
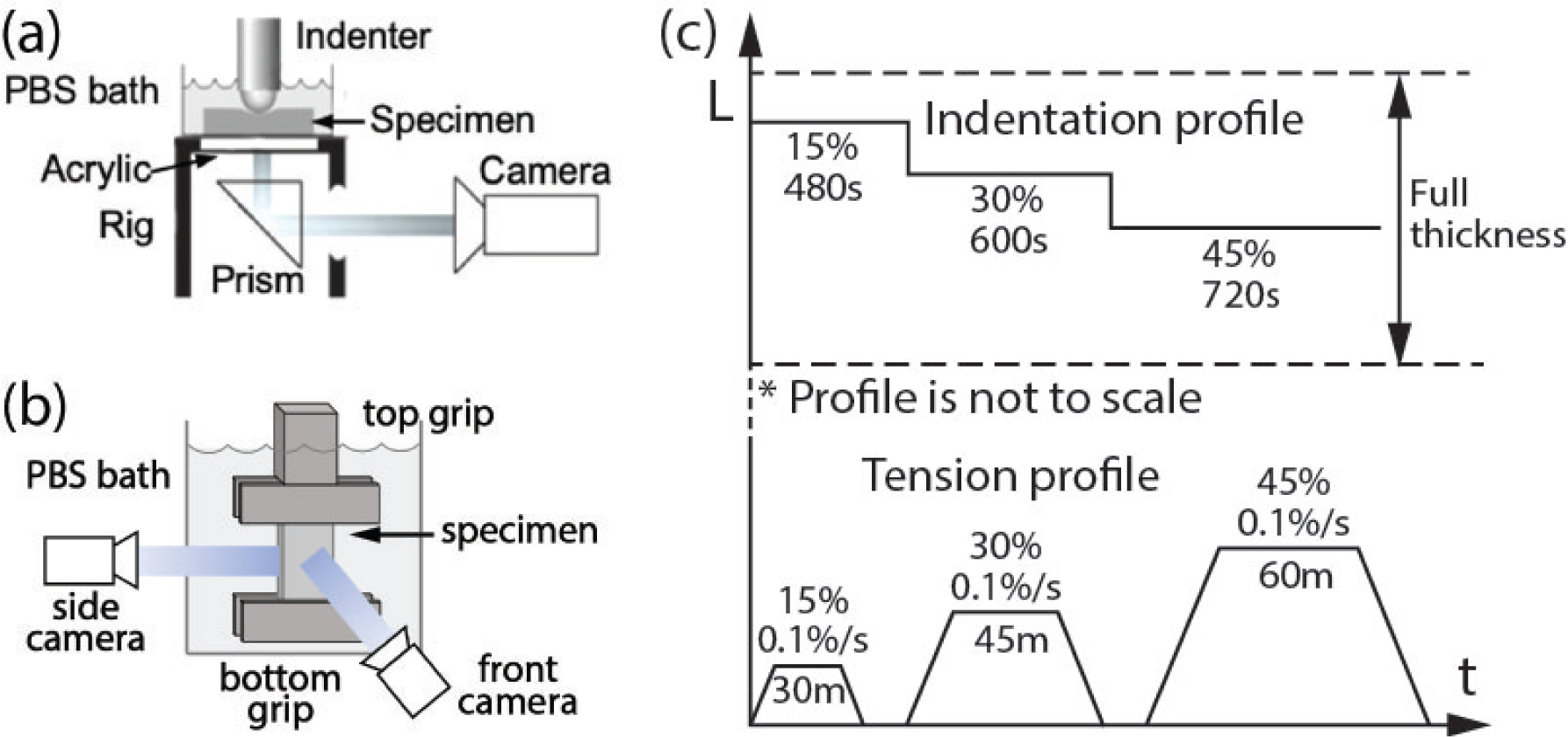
Mechanical test regime. The experiment illustrations for (a) indentation and (b) tension tests. (c) The protocol profiles for both tests. The strain rate for the indentation test was 1%*·*second*^−^*^1^.

### 2.3. Digital Image Data Processing

Digital image correlation (DIC) (Correlated Solutions, v6) and image segmentation (MATLAB) were performed following the previous workflow to measure and calculate the Green-Lagrange strain, stretch, and Cauchy stress.[12] For indentation, DIC was performed in a 2-mm diameter circle centered under the indenter (defined as the region of interest [ROI]). FEA shows an approximately uniform stress distribution within this circle (not shown here). For tension, because biological tissue is known to have non-homogeneous strain in the region of the grips, DIC was performed to characterize the 2D strain field.[21] A middle gauge region with an approximate uniform strain distribution was chosen as the region of interest (ROI) to ensure a uniaxial tensile condition. The finite element modeling used the strain from the DIC analysis instead of the G2G strain. Image segmentation was performed for the tension test recordings to extract three dimensions measuring the tensile specimen: length, width, and thickness. Stretch 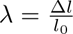 and Cauchy stress 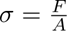 were subsequently calculated.

### 2.4. Equilibrium Material Behavior Characterization

To determine the equilibrium mechanical properties of the human uterus, inverse finite element analysis (IFEA) was performed based on the previously developed workflow described in Fang et al. [20]. Finite element (FE) models of the indentation and tension tests were built in the FE software FEBio Studio (Salt Lake City, Utah) with the setup illustrated in Fig. 3(a-b). The fiber-based constitutive law described in [19, 20] was adopted to model the material behavior. This constitutive model combines an isotropic compressible ground substance and an anisotropic extensible fiber network. The mechanical behavior of the ground substance is captured by shear and bulk terms with moduli *µ* and *κ*, respectively. The behavior of the collagen fiber distribution and the passive smooth muscle cells (SMC) is determined by material parameters such as the initial stiffness (*ξ*), locking stretch (*ζ*), and a fiber distribution parameter (*b*).

**Figure 3:**
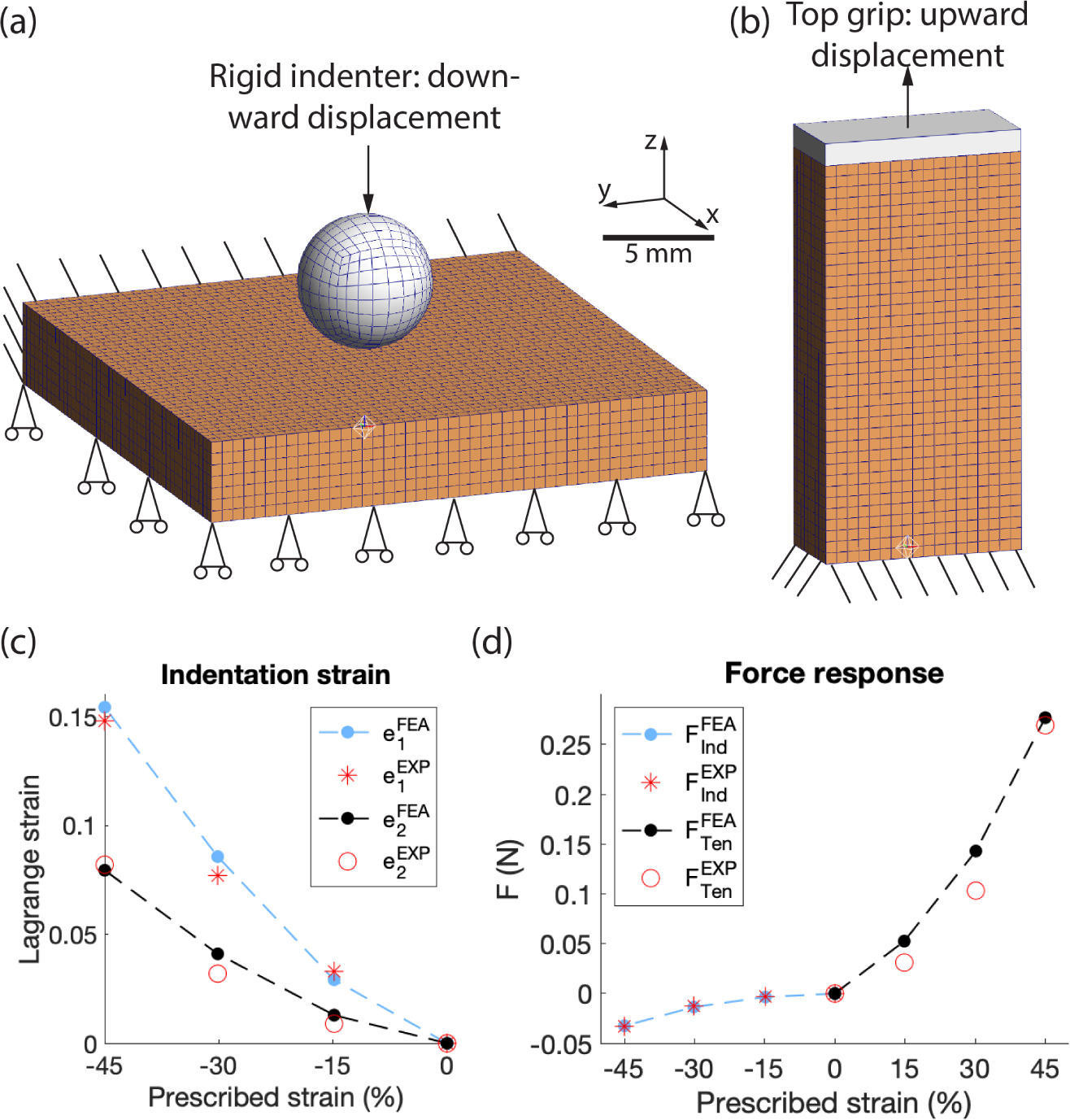
FEA models for mechanical tests and comparisons between FEA and experimental outcomes. (a) The indentation specimen was restrained in the z-direction at the bottom and fixed on one side to stabilize the geometry in space. (b) The tension specimen was fixed at its bottom to simulate the lower grips and the top grip was tied to the specimen top. Geometries were meshed using linear hexahedral elements with an edge length of 0.5 mm. (c-d) Lines and markers (asterisks and squares) represent the FEA and experimental responses, respectively. First (*e*_1_) and second (*e*_2_) principal strains and indentation (Ind) and tension (Ten) forces are plotted over displacements. Superscripts “FEA” and “EXP” represent variables of the finite element model and experiment, respectively.

In the IFEA, the constitutive model parameters were obtained by minimizing an objective function defined by the differences between the experimental and simulation results. The equilibrium force response from both spherical indentation and uniaxial tension tests, as well as the principal Lagrangian strains e_1_ and e_2_ from indentation, were incorporated into the calculation of the objective error function. A natural selection-based genetic algorithm (GA) implemented in MATLAB was used to minimize the objective function [22]. For this study, the population size was assigned as *n* = 4, the mutation rate *µ_m_* = 0.15, and a minimum of 40 generations were performed to reach an objective function value of less than 0.5 for all cases, ensuring a good fit and convergence of the solution.

Because the load response for some specimens did not reach equilibrium at the highest prescribed strain level of 45% in both the indentation and tension experiments, we obtained the equilibrium load by curve-fitting the experimental relaxation data using the following equation:

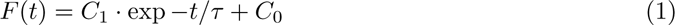

where *C*_0_, *C*_1_ and *τ* are constants. The new extrapolated equilibrium *F* = *C*_0_ was calculated for specimens whose difference between the extrapolated equilibrium load and the achieved load in the experiment was greater than 10%. Only the force data at times greater than 200 s and 100 s from the application of 45% strain were used for curve fitting in the indentation and tension experiments, respectively. Eq. 1 produced good fits for both indentation and tension force relaxation responses (R^2^ *>* 0.985 and 0.988 for Indentation and Tension, respectively).

In our previous human uterine study, optical coherence tomography (OCT) and curve fitting were used to characterize the fiber distribution of 27 uterine specimens (13 from one NP patient and 14 from one PG).[12] A von Mises distribution with a preferential direction *θ* and a concentration factor *b* was fitted to the directions of fiber bundles. Here, concentration is the reciprocal of the commonly used concept of dispersion. In this study, the fiber distributions of these 27 specimens were directly adopted. For the rest of the specimens, the first principal strain angle *γ* derived from the indentation DIC analysis was used to calculate the preferential fiber group direction *θ* with the assumption that *γ* is perpendicular to *θ*. This assumption is based on the anisotropy of a fibrous structure that the tensile modulus along the fiber is higher than one perpendicular to the fiber. Hence, fiber-embedded tissue is more likely to deform orthogonal to the preferential fiber direction under indentation. The concentration factor of the von Mises distribution was left as a fitting parameter in the IFEA.

### 2.5. Material Model Validation

Best-fit material models were validated against experimental data that were not directly used for IFEA. The same workflow as the NHP cervical study was adopted.[20] Briefly, the principal strain fields outside the ROI were validated for indentation, and the stretch ratios and Cauchy stress were validated for tension. Specifically, the first and second principal strains within a 4-mm diameter circle centered under the indenter were compared between the experiment and FEA. Experimental values were determined by DIC analysis and FEA values were simulation outputs. A normalized error map was generated using equation error 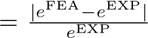 to quantify the comparison. Stretch ratios and Cauchy stress of the tension specimen were compared between the experiment and FEA. Experimental stretch ratios were calculated using dimensions derived from the image segmentation detailed in Sec. 2.3. Width and thickness (*λ*_1_ and *λ*_2_) were measured from the front and the side of the specimen. FEA values were direct simulation outputs.

### 2.6. Experimental Sensitivity Study

Experimental sensitivity studies were performed to assess the impact of experimental procedures on uterine tissue behaviors. Specifically, impacts caused by freeze-thaw cycles and the specimen topology were determined.

Due to the nature of tissue collection and the restrictions we faced in the operating room, the specimens went through two freeze-thaw cycles in total, from tissue collection to OCT scanning and mechanical testing. Many previous studies on soft tissues with similar components have found mixed results on whether and how freeze-thaw cycles affect the tissue’s mechanical properties.[23, 24] Therefore, comparison mechanical tests were performed on a different set of PG uterine tissue (16 C/S uteri) when they were fresh and frozen-thawed. These specimens were collected from women who underwent C/S at term for a non-pathological delivery, where an incision was made at the anterior lower uterine segment (approved by IRB). A small strip of the uterine sample was collected at the incision. Due to the nature of the collection, no orientation was recorded and the specimens varied in dimensions. Immediately after collection, the specimen was placed on wet ice and transferred over from the hospital to the laboratory and “freshly” tested within 2 hrs of collection. The indentation test was performed at the center of the specimen held in a PBS-filled petri dish. The protocol included three displacement-controlled steps with sufficient time in between for the specimen to relax and reach equilibrium. After the fresh test, the specimen was padded dry with paper towels and stored at *−*80 ℃ for more than 24 hours until the freeze-thaw test. During the freeze-thaw testing, the frozen specimen was thawed in PBS solution at room temperature for three hours. Then the same protocol was performed on the tissue at the same location. For both fresh and freeze-thaw testing, the force-time-displacement data were recorded and the equilibrium force responses at three levels were compared.

The human uterine tissue was dissected using a customized slicer designed to divide large samples into thinner even-surface specimens. In an effort to quantitatively determine whether the tissue surface is sufficiently smooth, a topology study was performed using spherical nanoindentation (Piuma, Optics11Life, Amsterdam, NE) on one NP and one PG slice. Using a 100 µm probe radius, two square areas of a 6 mm edge length were tested for each slice. The size of this square was equivalent to the maximum contact area during the indentation test. Within each area, 49 spots in a 7*×*7 equally-spaced mesh grid were tested, and the two areas were 5 mm apart (Fig. 4(a)). Three FEA models were then constructed, where the specimen had undulations (as tall as the largest detected height difference) on its top surface to simulate the unevenness, and the indenter tip was placed on top of, in the valley of, and on the ridge of the undulations (Fig. 4(b)). Due to the undulations on the surface, the geometry was meshed using linear tetrahedral elements with an edge length ranging from 0.5-0.7 mm. The same boundary conditions as the indentation test were implemented and all material parameters were set to be the average of all specimens. The indenter force and the first and second principal strains centered at the specimen bottom were compared between scenarios.

**Figure 4:**
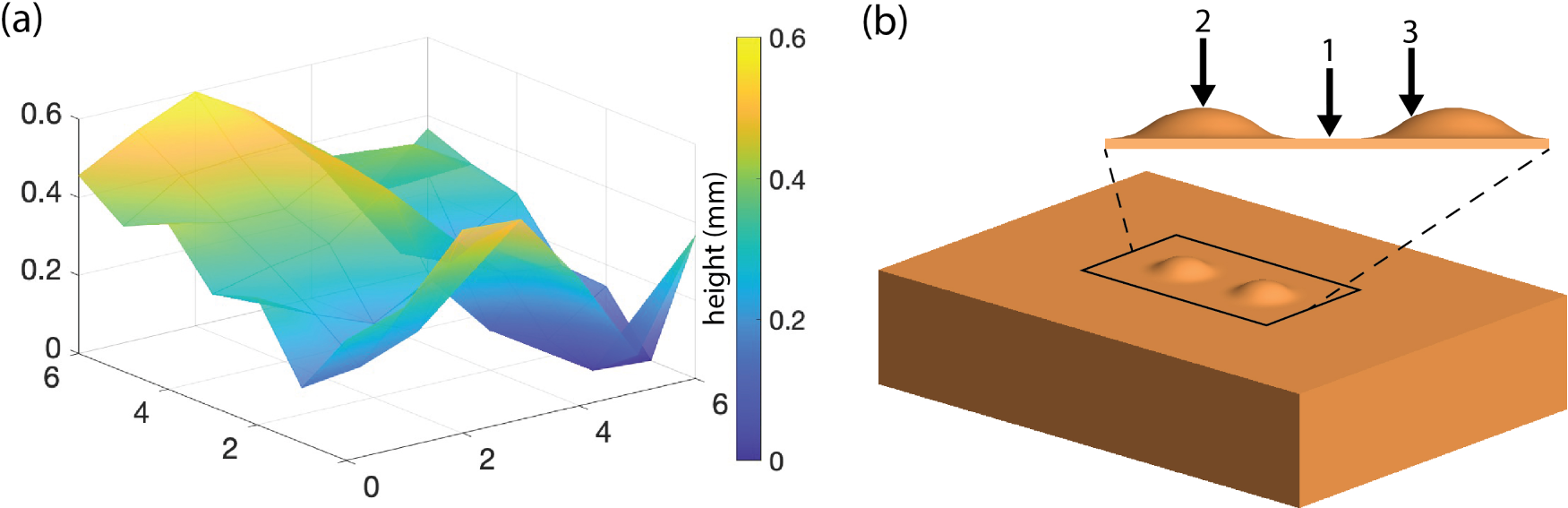
Sample topology study illustration. (a) A representative surface graph of one NP uterine sample shows that the top surface of the specimen has local variations in height. The largest height difference observed in this sample was around 0.6 mm. (b) An FEA model was constructed to simulate the undulating surface of the specimen. On the top of a specimen, two small undulations with a height of 0.6 mm were added 5 mm apart from each other. The locations of the indenter in three scenarios are indicated by the arrows numbered 1, 2, and 3 for between, on top of, and on the ridge of the undulations, respectively.

### 2.7. Statistical Analysis

D’Agostino and Pearson test was performed to examine the data normality. Welch’s t-test (*p*-value¡ 0.05) was then performed for normally distributed data, and Kolmogorov-Smirnov test was performed for non-normally distributed data.

## 3. Results

### 3.1. Mechanical Test Force, Strain, and Stress Response

Under indentation, the human uterus exhibits a force–relaxation response to a ramp–hold displacement protocol. Similar to our previous findings, both force and Lagrange strain responses at equilibrium are nonlinear with respect to prescribed strains (Fig. 5(a-b)).[12] Among all specimens, differences are observed between the first (*e*_1_) and second (*e*_2_) principal strains (Fig. 5(b)). The ratio distribution between these two strains of all specimens is 2.14 *±* 1.84.

**Figure 5:**
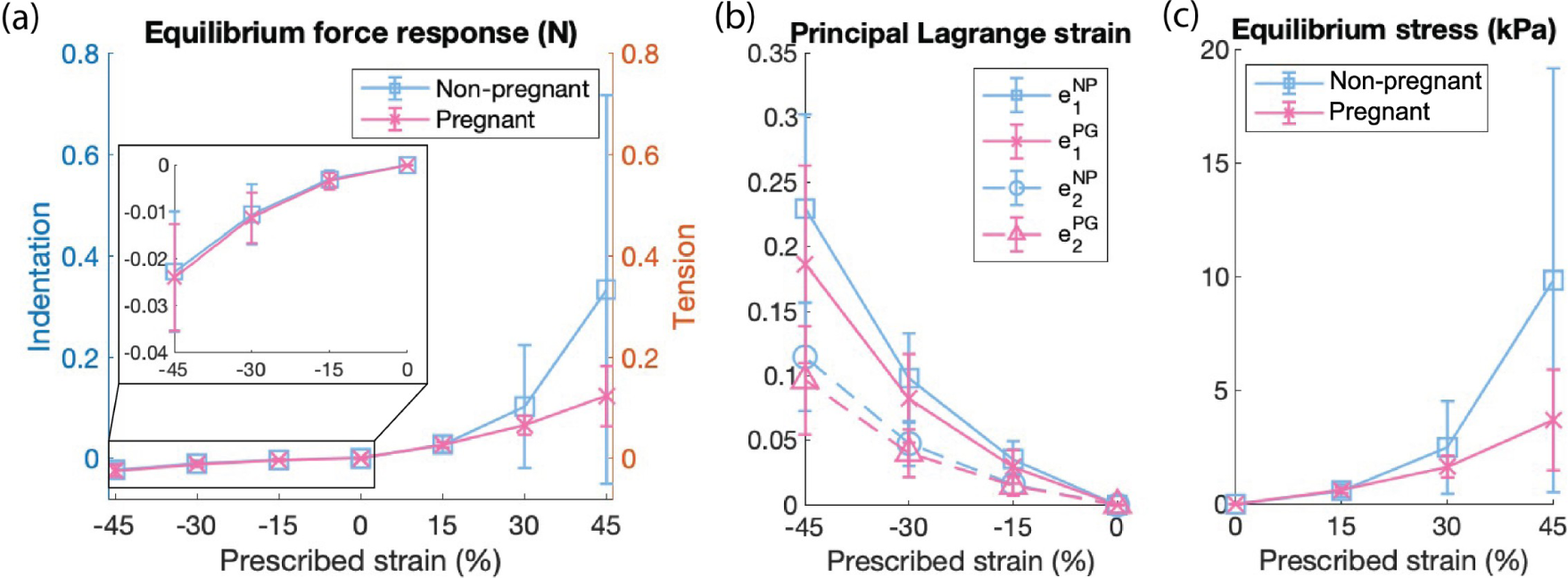
Equilibrium force and stress responses of the mechanical tests. (a) The force response, (b) the principal Lagrange strain response, and (c) the Cauchy stress response of the mechanical tests on all specimens are shown with mean *±* standard deviations. Negative values represent compressive variables in indentation while positive values represent tensile ones in tension. Indentation force responses are zoomed in within the black box in (a). Lines represent linearly-interpolated data.

Under tension, the human uterine tissue displays a J-shaped stress response to linear strains and a force–relaxation response to a load–hold displacement protocol (Fig. 6(a)). The Lagrange strain, force, and Cauchy stress response at equilibrium states are nonlinear to G2G strains (Fig. 6(b) and Fig. 5(a)(c)). The stress responses at the tensile equilibrium of the NP uterus are larger than the PG ones. For example, the equilibrium stress response distributions at the third strain level are 9.85 *±* 9.33 *kPa* and 3.70 *±* 2.22 *kPa* for NP and PG specimens, respectively (Fig. 5(c)). The strain fields of the human uterus during a uniaxial tension test are complex (Fig. 6(b)). Compared to the G2G strain levels of 15%, 30%, and 45%, the actual Lagrange strains across the specimen are non-homogeneous, larger than the G2G strain value in certain areas and smaller in others. The average values of the full field are similar to the prescribed strains. The edge of the specimen is excluded from the analysis to avoid geometric distortions from gripping. Within the analyzed area of this specific specimen, the top left corner and the middle gauge region show larger values whereas the lower quarter and the top right corner show smaller values. All specimen geometries and equilibrium force and strain data are available at the Columbia University Libraries’ Academic Commons (https://doi.org/10.7916/d8-r7mq-at21).

**Figure 6:**
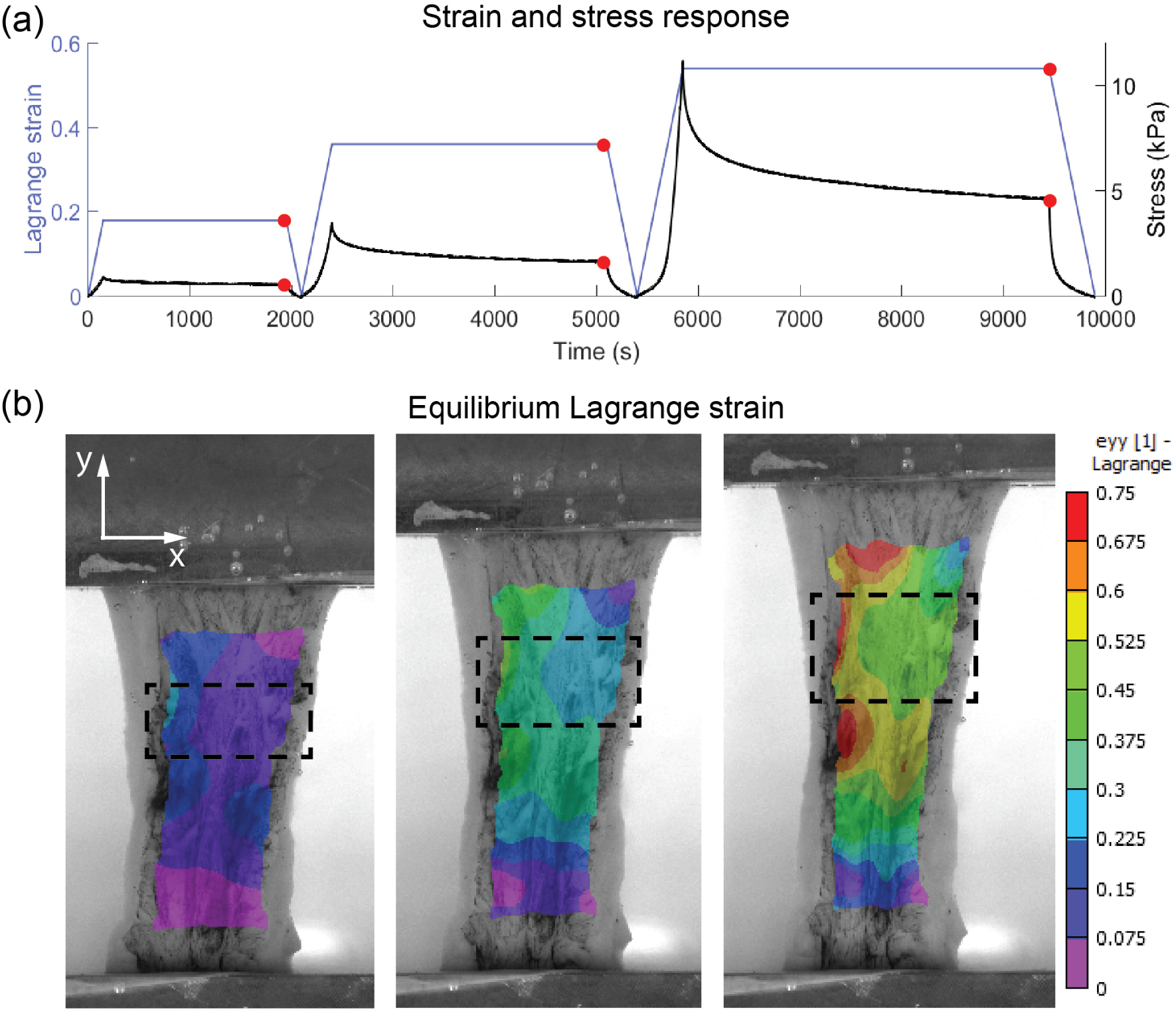
A representative stress–relaxation response and the Lagrange strain maps of a NP fundal tension specimen. (a) Three levels (15%, 30%, and 45% prescribed strain) of load–hold–unload cycles were applied and the Lagrange strain (in blue) of the ROI and stress (in black) responses of the specimen are plotted over time. Red dots mark the equilibrium states. (b) Equilibrium Lagrange strain maps show non-homogeneous distributions across the specimen in tension. Black dashed-line rectangles represent ROIs. The average strains across the entire analysis region are approximately equal to the prescribed strains.

### 3.2. Equilibrium Material Properties

The equilibrium mechanical properties of the human uterus for non-pregnancy and early third trimester are reported in this section (Fig. 7). The mean and standard deviation of every parameter for both gestation groups are listed in Table 2. The ground substance bulk modulus *κ*, fiber network locking stretch *ζ*, and initial stiffness *ξ* of all specimens are computed by the GA-based IFEA. Among four properties, (1) the bulk modulus *κ*, describing the ground substance compressibility, is larger in the average value of the NP tissue but shows no statistical significance; (2) the locking stretch *ζ*, describing the fiber network extensibility under loading, is significantly smaller for the NP tissue; (3) the initial stiffness *ξ* shows no difference between NP and PG specimens; and (4) the fiber concentration factor *b*, describing the fiber bundle group alignment, is significantly higher for the NP tissue, indicating a more aligned organization for fiber bundles. The value distributions of the locking stretch and the fiber concentration are larger for the NP tissue, while the value distributions of the bulk modulus are similar between NP and PG tissue. The bulk modulus essentially dictates the material’s resistance to volumetric change under hydrostatic pressure and it is the major contributor to the force response in indentation. The fiber concentration factor directly dictates the anisotropic behavior of the tissue.

**Figure 7:**
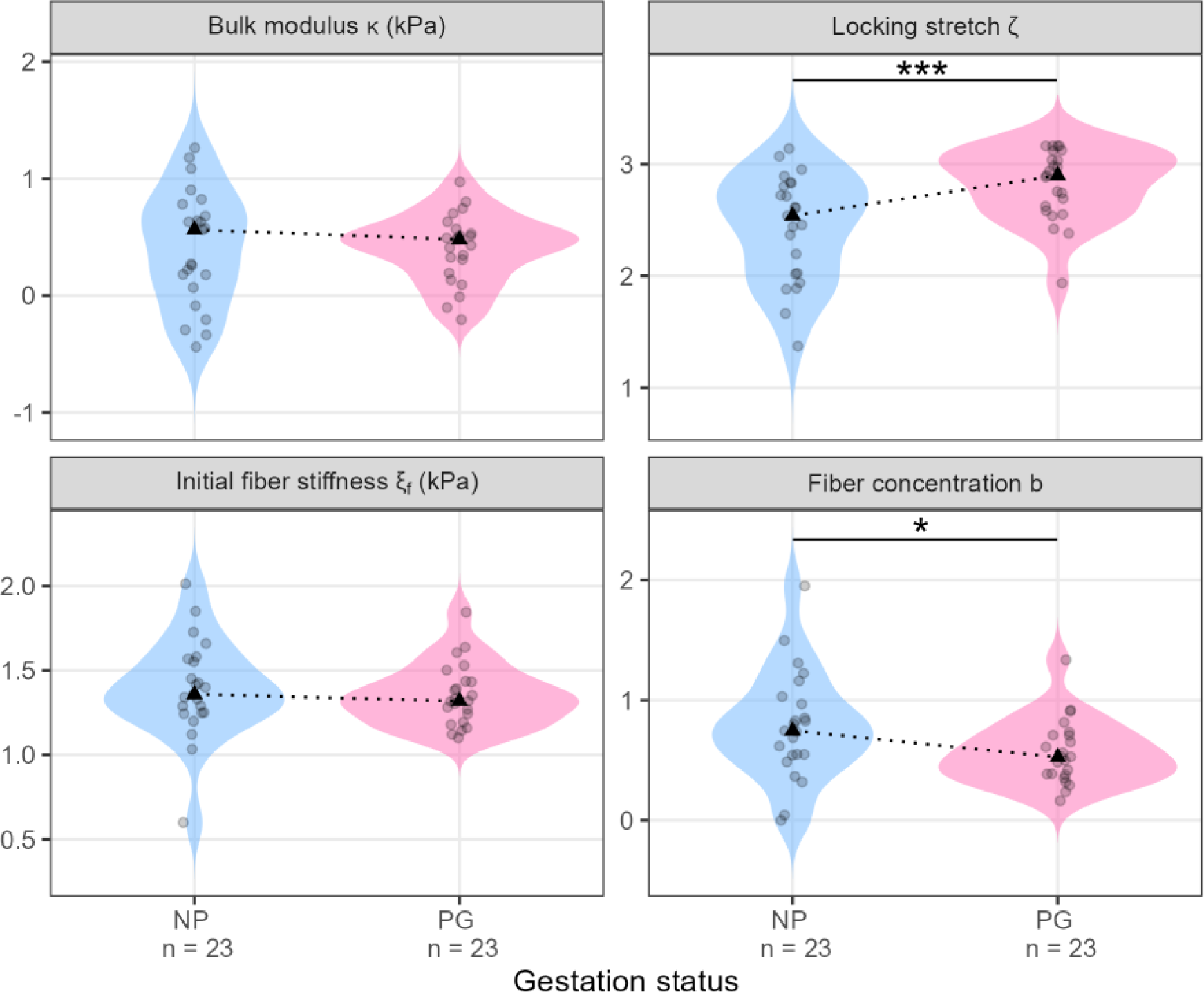
Mechanical properties of the human uterus. The bulk modulus *κ* and the initial fiber stiffness *ξ* are not significantly different between two time points. The locking stretch *ζ* of the PG tissue is higher than that of the NP tissue, indicating a more extensible fiber network under tensile loading. The fiber concentration factor *b* of the NP tissue is significantly higher than PG tissue, indicating more fiber bundles orient along the same direction, whereas PG fiber bundles are more dispersed. The varying widths of color-shaded areas indicate varying data distributions. Every transparent black dot represents one specimen. Black triangles represent medians connected by dotted black lines. Statistical significance: * *<* 0.05; ** *<* 0.01; *** *<* 0.005.

**Table 2:** Mechanical properties of the human uterus.

| Parameter | Mean value | Standard deviation |
| --- | --- | --- |
| $\kappa_{\text{NP}}$ [0.1, 100] kPa | 4.51 | 4.82 |
| $\kappa_{\text{PG}}$ [0.1, 100] kPa | 3.11 | 2.02 |
| $\zeta_{\text{NP}}$ [1.1, 3.33] | 2.43 | 0.48 |
| $\zeta_{\text{PG}}$ [1.1, 3.33] | 2.82 | 0.32 |
| $\xi_{\text{NP}}$ [0.1, 1000] kPa | 30.0 | 21.7 |
| $\xi_{\text{PG}}$ [0.1, 1000] kPa | 24.7 | 12.9 |
| $b_{\text{NP}}$ [0, 1] | 0.79 | 0.47 |
| $b_{\text{PG}}$ [0, 1] | 0.56 | 0.27 |

Subscripts “NP” and “PG” represent data from NP ( $N = 23$ ) and PG ( $N = 23$ ) specimens, respectively; Values in square brackets represent the search space of each parameter in IFEA based on preliminary results.

### 3.3. Material Model Validation

In general, the material model captures tissue behaviors well under both indentation and tension as shown by a single NP uterine sample (Fig. 8) In indentation, the model closely matches the experiment in terms of the magnitude, pattern, and direction of principal Lagrange strains of the bottom surface (Fig. 8(a-c)). In this representative specimen at the third equilibrium (45% prescribed strain) level, the first principal strain has a maximum of about 0.25 around the center and an elliptical distribution pattern with a nearly horizontal major axis. The directions of the strain close to the center point toward the positive y-direction. The second principal strain has a maximum of 0.15 around the center and an elliptical distribution pattern with a nearly vertical major axis. The directions of the strain close to the center point toward the negative x-direction. These features are captured well by the FEA model (Fig. 8(b)). When comparing the principal strain outside the ROI for IFEA, the strains of the experiment and the material model match well in terms of magnitude, shown by the small errors (Fig. 8(c)). However, areas further away from the center exhibit larger errors. In tension, the model matches the experiment well with regard to the Cauchy stress and the orthogonal stretch ratios (Fig. 8(d-f)). In this representative specimen, the material model captures the magnitude and the nonlinearity of the Cauchy stress. The sample deformations of the material model also closely match the experiment from both the front and the side.

**Figure 8:**
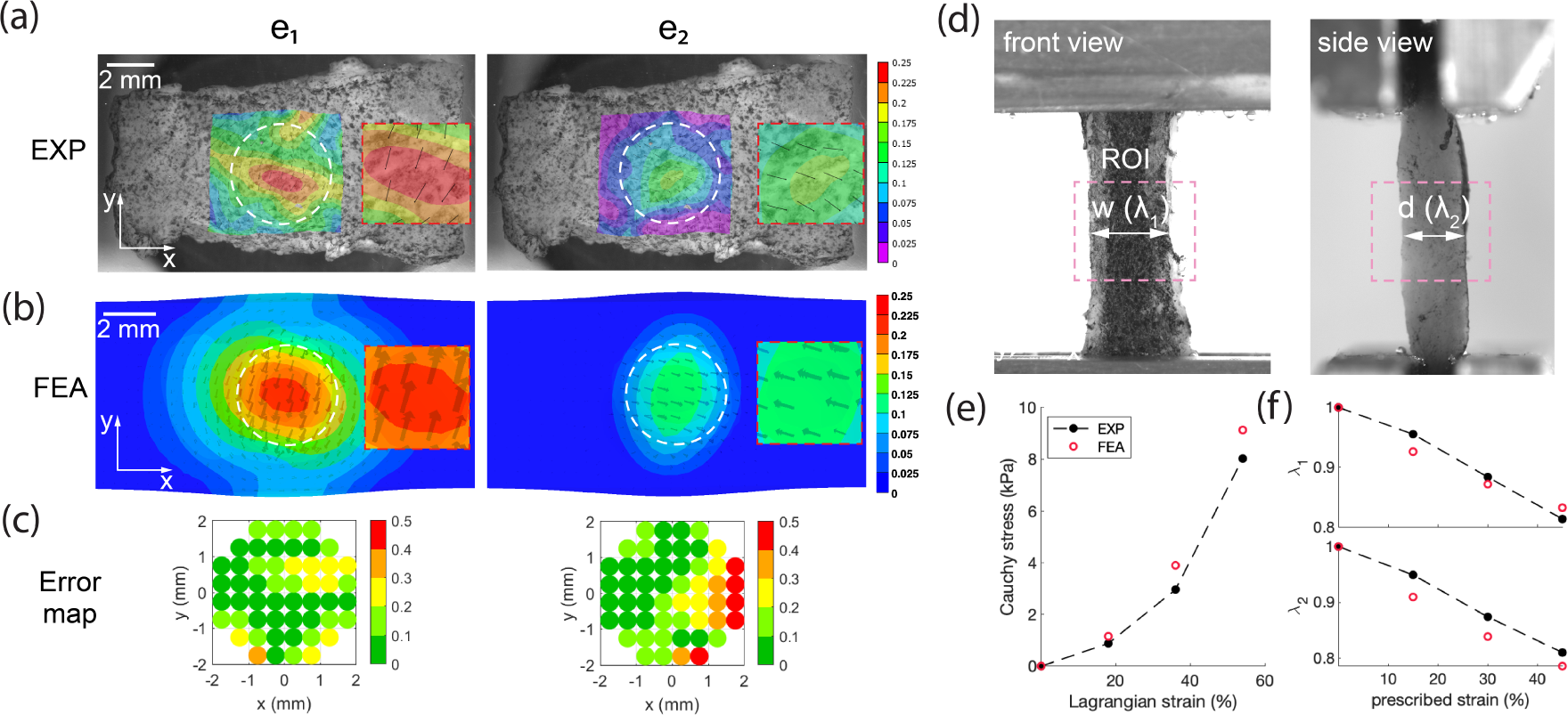
Material model validation. The first (*e*_1_) and second (*e*_2_) principal Lagrange strains of a representative indentation specimen bottom surface are compared between (a) the experiment and (b) the FEA model. The black lines in (a) represent the strain directions pointing away from the circle ends. The black arrows in (b) represent the strain directions pointing toward the arrow ends. Red dashed-line rectangles enclose magnified views of the strain directions. (c) Error maps of a 4-mm diameter circle are centered around the indenting location (0, 0). Errors greater than 0.5 are set to 0.5 for a more refined presentation. (d) The width (w) and thickness (t) and corresponding stretch ratios *λ*_1_ and *λ*_2_ of a representative tension specimen are measured from orthogonal images. The pink dashed-line rectangles enclose the ROI for dimension calculation. (e) The Cauchy stress at equilibrium is plotted with respect to the front surface Lagrange strain. (f) The stretch ratios at equilibrium are plotted with respect to prescribed strains. Black markers represent experiments, and red ones represent the model. Dashed lines between circles represent data by linear interpolation. Data are from a NP specimen.

### 3.4. Experimental Sensitivity

After one freeze-thaw cycle, the tissue behavior under indentation is similar to fresh tissues (Fig. 9). The raw force responses at equilibrium are similar between these two scenarios. No systematic alterations of tissue properties were observed as the median changes in percentage center around zero (Fig. 9(b)). Although the dimensions are not consistent across all specimens due to tissue collection restrictions and the raw force response is not a normalized variable, the comparison is performed within each sample that underwent the same testing protocol before and after freezing. For specimen topology effect, scenarios 1 to 3 are when the indenter is placed on top of, in the valley of (between), and on the ridge of undulations. For the principal strains, scenario 2 shows higher values than scenario 3 (followed by scenario 1) at the first and second strain levels. At the last strain level, all three scenarios exhibit similar principal strain values. For the indenter force response, scenario 2 shows a higher value than scenario 3, followed by scenario 1, and this difference exists for all three strain levels. However, all differences at the third strain level are less than 5% between scenarios.

**Figure 9:**
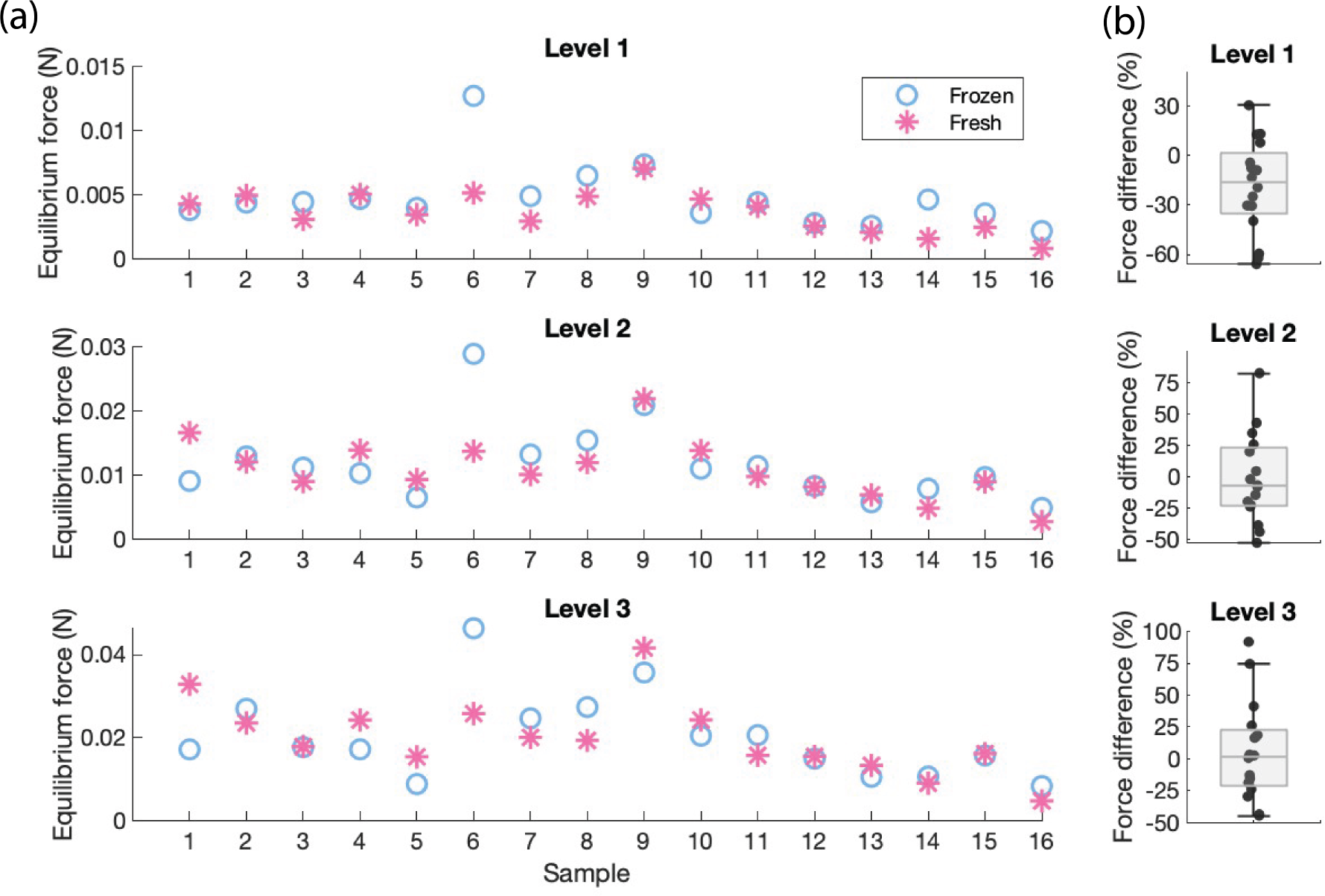
Freeze-thaw effects on uterine tissue behaviors. (a) Equilibrium force responses of three indentation levels between fresh (pink) and frozen-thawed (blue) tissues. Levels 1-3 correspond to prescribed strains of 10-30%. 16 samples are tested and indicated by numbers on the x-axis. (b) The difference in force responses is divided by the fresh tissue response for normalization. Every black dot represents one specimen. The middle horizontal lines of the boxes denote the medians.

## 4. Discussion

### 4.1. Human Uterus Material Behaviors

The material behaviors of the human uterine tissue under both compressive and tensile loads are time-dependent, non-homogeneous, and nonlinear (Fig. 6(a), [12]). Further, similar to human and NHP cervical tissue, the human uterus is anisotropic under indentation (Fig. 5(b)) and exhibits tension-compression asymmetry,.[22, 20, 19]

#### 4.1.1. Time-dependency

The human uterus displays a force–relaxation response to a load–hold displacement (Fig. 6(a), [12]). The time dependency of the human uterus is likely a product of viscoelastic and poroelastic mechanisms which can be attributed to the uncrimping and rearrangement of the fiber network and the escape of fluid through the pores of the tissue, respectively. Parallel work by our group has characterized the time-dependent behavior of NP and PG uterine tissue from an overlapping patient cohort using nanoindentation.[14] Analysis of the data with a combined poroelastic-viscoelastic model reveals that the innermost uterine layer, known as the endometrium-decidua, exhibits the greatest alterations in time-dependent material properties in pregnancy as evidenced by changes in viscoelastic ratio and diffusivity parameters.[14] No change in the time-dependent properties of the myometrium is observed at this micrometer length scale.[14] Future work is still needed to determine the degree to which poroelastic and viscoelastic mechanisms contribute to the overall time-dependent behavior of the uterus. [25]

#### 4.1.2. Non-homogeneity

The tensile Lagrangian strain fields are non-homogeneous across the uterine specimen. Several concurrent mechanisms could contribute to this phenomenon, including collagen fiber alignment, fiber group interactions, and the existence of different tissue structural components within one area. Previous work from our lab which characterized the human uterine fiber orientation within a small area found that multiple fiber families with different preferential directions exist.[12] The actual mechanism behind this observation remains an open question due to the lack of comprehensive histological and microscopic inferences.

#### 4.1.3. Nonlinearity

The human uterus displays a nonlinear force, strain, and stress response to linear prescribed strains. This nonlinearity manifests at two scales, the instantaneous/temporal behavior at every displacement cycle (Fig. 6(a)) and the equilibrium behavior across multiple strain levels ((Fig. 5). At the instantaneous state, the tensile stress response resembles a J-shaped curve consisting of two different segments, a low-slope linear region followed by a transition to a higher-sloped linear one. This indicates at least two different mechanisms behind the tissue stress responses. Fibrous biological tissues are known to have similar behavior [26]. The contribution of the ground substance and elastic fibers are associated with the small-strain regime, whereas the engagement of the collagen fibers is found at larger strains [26]. Across multiple equilibrium states, both indentation and tension responses exhibit a curve with an increasing slope. The rate of increase in the force response is larger for tension than indentation. Wavy fibers mainly hold tension and have a more pronounced effect on mechanical properties in the tension tests. Indentation applies a complex stress condition, and the tensile properties of the fibers engage due to Poisson’s effect at large compressive stresses.

#### 4.1.4. Anisotropy

Under indentation for a given ROI, the first and second principal strain fields differ significantly, indicating an anisotropic tissue behavior. This anisotropy is dictated by the fiber alignment described by the fiber group’s preferential orientation and the fiber bundles’ dispersion. Our results show a higher fiber concentration factor is associated with a larger anisotropy. When fiber bundles are less dispersed, this group of fibers exhibits greater direction-dependent behaviors. Under tension, the Lagrange strain fields are non-homogeneous across the specimen and the strain patterns are inconsistent across different samples. The non-homogeneity of the strain pattern is most likely the result of, although not limited to, the directional fiber orientations, another manifestation of anisotropy. Collagen fibers are known to have higher stiffness along their length versus orthogonally, and hence, deform less when loading aligns with the length dimension. The strain field is complex when multiple fiber groups interact within a single tissue by interweaving or crossing. The pattern of the strain field is also dynamic over time as observed between different equilibrium G2G strain levels (Fig. 6(b)). The fiber network movement likely causes this changing pattern. We hypothesize that the fibers rearrange themselves under loading by uncrimping and untangling to resist deformation and avoid damage, thus causing the changing strain patterns.

#### 4.1.5. Tension-Compression Asymmetry

A significant asymmetry was observed when comparing uterine material behaviors between indentation and tension. The tensile Cauchy stress is an order of magnitude higher than the indentation stress at the maximum. A combination of the fibers’ robe-like structure and the ground substance’s low stiffness dictates the primary contribution to this phenomenon. Fibers cannot engage under compressive loading due to their rope-like structure and hence, do not contribute significantly to the force response under indentation. Instead, the ground substance engages in compressive resistance, and its low stiffness contributes to the lower force response. Lastly, although uterine tissue exhibits time-dependent properties under both indentation and tension, the primary deformation mechanisms are thought to differ between these conditions. Under tension, as the fiber network rearranges its alignment, stress relaxation is thought to be dominated by the viscoelastic nature of the fiber network. In compression, the ground substance’s volumetric stiffness resists hydrostatic pressures. Therefore, under indentation, the measured force relaxation is thought to be dominated by poroelastic mechanisms draining the pressure from interstitial pores. Additional experiments and modeling efforts are warranted to characterize these time-dependent features of the material.

### 4.2. Human Uterus Equilibrium Mechanical Properties

Overall, bulk modulus *κ* of the ground substance is 4.39 *±* 4.33 kPa for the NP uterus and 3.93 *±* 3.71 kPa for the PG uterus, larger than the human cervix (1.40 *±* 1.40 kPa for NP and 0.26 *±* 0.20 kPa for PG) but is at the same order of magnitude. This difference indicates that the human uterus has a lower compressibility level than the cervix.[19] Though not statistically significant, the ground substance of the NP uterus is, on average, less compressible than the PG tissue. This trend is the same as the human uterus characterized using a neo-Hookean-based model under only indentation[12]. The locking stretch *ζ* of the PG tissue (2.74 *±* 0.33) is significantly larger than the NP uterus (2.21 *±* 0.40), indicating a more extensible tissue property for the PG uterus. This difference in extensibility is observed in the tension test, where PG tissue can extend more than the NP tissue when subjected to the same amount of force. The initial stiffness *ξ* describes the small-strain stiffness and is almost identical between the NP and PG tissue, indicating the behavior of the uterine tissue under small strain does not change over pregnancy. Previous work done in our lab characterized the human uterus stiffness under small strains using an isotropic material model and found a similar trend[12]. The fiber concentration *b* is the reciprocal of fiber dispersion and the PG fiber network is more dispersed than the NP one. A higher dispersion facilitates the fiber network’s extensibility under tensile loading; this change is also observed in the nonhuman primate cervix.[20] Finally, although both direct stiffness measurements, bulk modulus and the initial fiber stiffness, do not change significantly over pregnancy, the overall stiffness of the uterus is a combined contribution from *κ*, *ζ*, and *ξ*. Therefore, the increase in locking stretch, which is dominant under tension, indicates that the human uterus is less stiff during late pregnancy than in nonpregnancy. In other words, there is no stiffness change in the initial loading, but there is an overall difference when accounting for larger strains (*>* 0.3).

#### 4.2.1. Multiscale Mechanical Properties

To adequately perform essential reproductive functions, the uterus exhibits a unique hierarchy of structural components, spanning from micro- to milli-meter length scales. The relative contribution of different tissue components varies depending on the loading condition. Macro-scale mechanical testing used in this study measures the combined contribution of smooth muscle fascicles and collagen fibers under tension and compression at high strains. In contrast, micro-mechanical testing probes on the scale of individual smooth muscle cells and collagen fibers. Yet, despite these differences in length scales, there is a striking similarity in the data trends observed for the myometrium, as shown by a parallel study conducted in our lab[14]. Interestingly, no change in stiffness is observed for the myometrium in pregnancy at low strains for both micro- and macro-scale testing.[14] Further, both testing modalities yield stiffness values for the myometrium within the same order of magnitude (0.1 - 1 kPa). Differences in the mechanical behavior of pregnant uterine tissue only emerge under large deformations for strains above 0.15, as demonstrated by increased tissue extensibility.

### 4.3. Constitutive Modeling Approach

A constitutive model previously developed for the human cervix is adopted in this work to capture the material behaviors of the NP and PG human uterus [19]. Compared to the cervix, the uterus has a larger proportion of smooth muscle cells (SMCs) to collagen content[27]. Despite these basic compositional differences, the approach of adopting the same material model is supported by the following considerations. First, mechanically, SMCs play an important role in active material behaviors instead of passive equilibrium behaviors, as they contribute to the force response by contracting. The passive material response to loading of the uterus shares similar patterns with that of the cervix, a material mostly composed of collagens and doesn’t contract. Second, also mechanically, the stiffness modulus of the muscle-collagen fiber composite of the skeletal muscle tissue was found to be five times the muscle fiber modulus alone, indicating collagens are the primary passive load-bearing component in the tissue.[28] Third, biologically, SMCs are surrounded by a collagenous membrane called endomysium, which separates and groups smooth muscle fibers, resulting in larger fiber bundles of the same alignment. Consequently, the combination of smooth muscle fibers and collagen fibers in the uterus was treated together as one fiber network, a condition similar to the cervix and suitable for the previously developed material model.

### 4.4. Experimental and Computational Error Analysis

Inherent experimental errors include the load cell tolerance (0.005 N), the displacement resolution of the Instron (8 *µ*m), and the buoyancy force on the indenter (*F_b_ < ρ*gV *<* 0.001 N) and the tensile grips (*<* 0.009 N). As the maximum difference in all three variables under three scenarios of sample topology is less than 5% (Fig. 10) and the samples do not have larger variations in the height of the top surface, the effect of sample topology is negligible. Inherent computational errors introduced by the DIC process and fixing the far-end specimen side surface in FEA are both found to be less than 2%.[20, 12] GA-based IFEA was examined the same way as our previous study to ensure the global optimum is found.[29]

**Figure 10:**
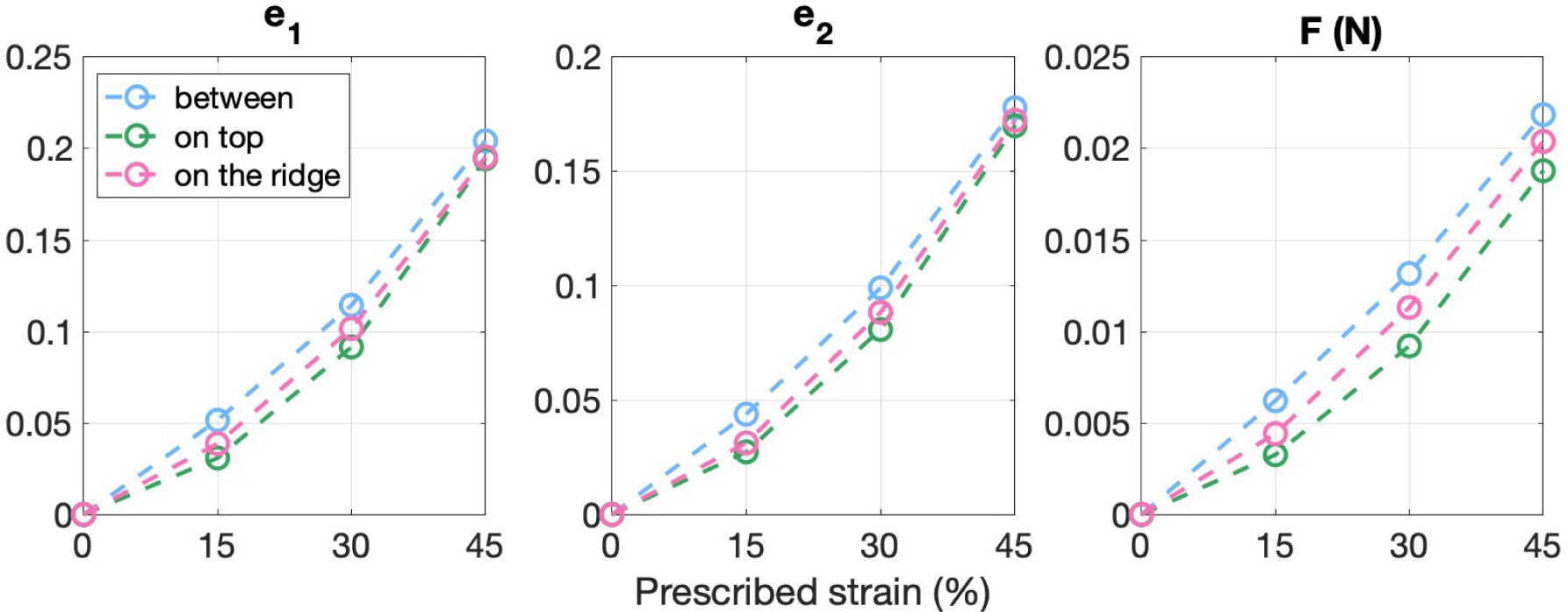
Tissue behavior variations by topology in FEA. The first and second (*e*_1_ and *e*_2_) principal Lagrange strains of the specimen bottom and the force response (F) recorded by the indenter were compared between the three scenarios. Data were recorded at three indentation strains and are linearly interpolated between data points.

### 4.5. Limitations

The validation study shows that the fitted material model captures principal strains effectively within the 4-mm diameter circle around the indenter, but less accurately near the edge of this circle (Fig. 8(c)). This is expected for the following reasons: first, the fiber distribution outside the original ROI (2-mm diameter circle) is different and not characterized in the IFEA; second, the stress closer to the edge of the circle differs more from the center under spherical indentation. Although smooth muscle fibers and collagen fibers share many similarities in biological structures and mechanical functions, the assumption of treating these two as one fiber network may be an oversimplification. However, our current approach is taken as there is no further knowledge of the interactions between these two fibers. For obvious reasons, specimens collected from a hysterectomy are often pathological (Table 1), which may contribute to alterations in normal mechanical properties.[12] Ethnicity is demonstrated to affect tissue behaviors elsewhere,[30] but the effect is not explored here. Biological tissues behave differently under differing loading conditions, and hence, the reported material properties should only be applied to similar strain regimes. Only for patients NP1 and PG1, the anterior and posterior myometrium tissue were studied for reasons explained in sec. 2.1. Fundal myometrium tissue was measured for the rest of the study participants. As the mechanical properties of the human uterus were found to be inherently heterogeneous across anatomic regions and tissue layers, the reported results should be applied with consideration.

## 5. Conclusions

In this study, the equilibrium mechanical properties of the human uterus under indentation and tension are characterized using a single set of material parameters. The fiber architecture (directionality and dispersion) was characterized using OCT and incorporated into the material model to describe tissue anisotropy. The oriented fiber network dictates tissue strains under both indentation and tension. Although spherical indentation results in a complex stress distribution, it reveals the directional material behavior and preserves specimen integrity to enable follow-up tension tests. While indentation testing primarily assesses the response of the ground substance, tension testing fully engages the fiber network for mechanical characterization. A 3-dimensional anisotropic constitutive model is adapted from a human cervical study to assess to mechanical properties of the NP and PG uterus, given the similarities of the two tissue types. The model is optimized using inverse finite element analysis to best fit the mechanical tests and validated against the experiments. The material parameters correlate with the tissue microstructure and can capture the structure–function relationship. The PG human uterus in the third trimester is found to be (1) less stiff, (2) more extensible, and (3) more dispersed in its fiber network compared to nonpregnancy. This study is the first to characterize both compressive and tensile equilibrium behaviors of the NP and PG human uterus using one set of parameters. The mechanical characterization of the human uterus is fundamental to guiding other research in biomechanics, mechanobiology, biomaterials, and tissue engineering, as well as informing the development of novel therapeutic approaches for obstetric and gynecologic conditions.

## Acknowledgement

The authors thank Dr. Fady Khoury-Collado from Columbia University Irving Medical Center Department of Obstetrics and Gynecology and Dr. George Gallos from Walnut Creek Medical Center Department of Anesthesiology for uterine tissue specimens. Research reported in this publication was supported by the Eunice Kennedy Shriver National Institute of Child Health & Human Development under Awards R01HD091153 and R01HD072077 to KMM and the Iris Fund. The content is solely the responsibility of the authors and does not necessarily represent the official views of the National Institutes of Health.

## Notes

### Competing Interest Statement

The authors have declared no competing interest.

## References

[1] P. Sokolowski, F. Saison, W. Giles, S. McGrath, D. Smith, J. Smith, R. Smith, Human Uterine Wall Tension Trajectories and the Onset of Parturition, PLoS ONE 5 (6) (2010) e11037. doi:10.1371/journal.pone.0011037. URL https://dx.plos.org/10.1371/journal.pone.0011037

[2] M. Ono, T. Maruyama, H. Masuda, T. Kajitani, T. Nagashima, T. Arase, M. Ito, K. Ohta, H. Uchida, H. Asada, Y. Yoshimura, H. Okano, Y. Matsuzaki, Side population in human uterine myometrium displays phenotypic and functional characteristics of myometrial stem cells, Proceedings of the National Academy of Sciences 104 (47) (2007) 18700–18705. doi: 10.1073/pnas.0704472104. URL https://pnas.org/doi/full/10.1073/pnas.0704472104

[3] J. L. Cook, D. B. Zaragoza, D. H. Sung, D. M. Olson, Expression of myometrial activation and stimulation genes in a mouse model of preterm labor: Myometrial activation, stimulation, and preterm labor, Endocrinology 141 (5) (2000) 1718–1728. doi:10.1210/endo.141.5.7474.

[4] Born too soon: decade of action on preterm birth. URL https://www.who.int/publications-detail-redirect/9789240073890

[5] Born too soon: The global action report on preterm birth. URL https://www.who.int/publications-detail-redirect/9789241503433

[6] World Health Organization: Preterm birth. URL https://www.who.int/news-room/fact-sheets/detail/preterm-birth

[7] J. Alexander, Prolonged pregnancy: induction of labor and cesarean births, Obstetrics & Gynecology 97 (6) (2001) 911–915. doi:10.1016/S0029-7844(01)01354-0. URL http://linkinghub.elsevier.com/retrieve/pii/S0029784401013540

[8] R. L. Goldenberg, J. F. Culhane, J. D. Iams, R. Romero, Epidemiology and causes of preterm birth, The Lancet 371 (9606) (2008) 75–84. doi:10.1016/S0140-6736(08)60074-4. URL https://linkinghub.elsevier.com/retrieve/pii/S0140673608600744

[9] A. Many, L. Hill, N. Lazebnik, J. Martin, The association between polyhydramnios and preterm delivery, Obstetrics & Gynecology 86 (3) (1995) 389–391. doi:10.1016/0029-7844(95)00179-U. URL http://linkinghub.elsevier.com/retrieve/pii/002978449500179U

[10] J. T. Conrad, W. L. Johnson, W. K. Kuhn, C. A. Hunter Jr, Passive stretch relationships in human uterine muscle, American Journal of Obstetrics and Gynecology 96 (8) (1966) 1055–1059.

[11] C. Wood, The expansile behaviour of the human uterus, BJOG: An International Journal of Obstetrics & Gynaecology 71 (4) (1964) 615–620.

[12] S. Fang, J. McLean, L. Shi, J.-S. Y. Vink, C. P. Hendon, K. M. Myers, Anisotropic Mechanical Properties of the Human Uterus Measured by Spherical Indentation, Annals of Biomedical Engineering 49 (8) (2021) 1923–1942. doi:10.1007/s10439-021-02769-0. URL https://link.springer.com/10.1007/s10439-021-02769-0

[13] S. Weiss, T. Jaermann, P. Schmid, P. Staempfli, P. Boesiger, P. Niederer, R. Caduff, M. Bajka, Three-dimensional fiber architecture of the nonpregnant human uterus determined ex vivo using magnetic resonance diffusion tensor imaging, Anatomical Record - Part A Discoveries in Molecular, Cellular, and Evolutionary Biology 288 (1) (2006) 84–90. doi:10.1002/ar.a. 20274.

[14] D. M. Fodera, S. R. Russell, J. L. Jackson, S. Fang, X. Chen, J. Vink, M. L. Oyen, K. M. Myers, Material properties of nonpregnant and pregnant human uterine layers, Journal of the Mechanical Behavior of Biomedical Materials 151 (2024) 106348. doi:10.1016/j.jmbbm.2023.106348.

[15] K. Dewilde, M. Vanthienen, D. Van Schoubroeck, W. Froyman, D. Timmerman, T. Van den Bosch, Elastography in ultrasound assessment of the uterus, Journal of Endometriosis and Uterine Disorders (2023) 100014.

[16] J. McLean, S. Fang, G. Gallos, K. Myers, C. Hendon, 3-D collagen fiber mapping and tractography ofhuman uterine tissue using OCT, Biomedical Optics Express 11 (10) (2020) 5518–5541. doi:10.1364/boe.397041.

[17] K. M. Myers, D. Elad, Biomechanics of the human uterus, Wiley Interdisciplinary Reviews: Systems Biology and Medicine 9 (5) (2017) 1–20. doi:10.1002/wsbm.1388.

[18] J. Woessner, T. Brewer, FORMATION AND BREAKDOWN OF COLLAGEN AND ELASTIN IN THE HUMAN UTERUS DURING PREGNANCY AND POST-PARTUM INVOLUTION, Biochemical Journal 89 (1) (1963) 75–82. doi:10.1042/bj0890075. URL https://portlandpress.com/biochemj/article/89/1/75/52979/ FORMATION-AND-BREAKDOWN-OF-COLLAGEN-AND-ELASTIN-IN

[19] L. Shi, L. Hu, N. Lee, S. Fang, K. Myers, Three-dimensional anisotropic hyperelastic constitutive model describing the mechanical response of human and mouse cervix, Acta Biomaterialia 150 (2022) 277–294. doi:10.1016/j.actbio.2022.07.062. URL https://linkinghub.elsevier.com/retrieve/pii/S1742706122004640

[20] S. Fang, L. Shi, J.-S. Y. Vink, H. Feltovich, T. J. Hall, K. M. Myers, Equilibrium Mechanical Properties of the Nonhuman Primate Cervix, Journal of Biomechanical Engineering 146 (8) (2024) 081001. arXiv:https://asmedigitalcollection.asme.org/biomechanical/article-pdf/146/8/081001/7321760/bio\_146\_08\_081001.pdf, doi:10.1115/1.4064558. URL 10.1115/1.4064558

[21] K. F. Mallett, E. M. Arruda, Digital image correlation-aided mechanical characterization of the anteromedial and posterolateral bundles of the anterior cruciate ligament, Acta Biomaterialia 56 (2017) 44–57. doi:10.1016/j.actbio.2017.03.045. URL https://linkinghub.elsevier.com/retrieve/pii/S1742706117302179

[22] L. Shi, W. Yao, Y. Gan, L. Y. Zhao, W. Eugene McKee, J. Vink, R. J. Wapner, C. P. Hendon, K. Myers, Anisotropic Material Characterization of Human Cervix Tissue Based on Indentation and Inverse Finite Element Analysis, Journal of biomechanical engineering 141 (9). doi:10.1115/1.4043977.

[23] M. Adham, J.-P. Gournier, J.-P. Favre, E. De La Roche, C. Ducerf, J. Baulieux, X. Barral, M. Pouyet, Mechanical characteristics of fresh and frozen human descending thoracic aorta, Journal of Surgical Research 64 (1) (1996) 32–34. doi:10.1006/jsre.1996.0302. URL https://linkinghub.elsevier.com/retrieve/pii/S0022480496903029

[24] J. Delgadillo, S. Delorme, R. El-Ayoubi, R. Diraddo, S. Hatzikiriakos, Effect of freezing on the passive mechanical properties of arterial samples, J. Biomedical Science and Engineering 337088 (2010) 645–652. doi:10.4236/jbise.2010.37088.

[25] K. Yoshida, C. Jayyosi, N. Lee, M. Mahendroo, K. M. Myers, Mechanics of cervical remodelling: insights from rodent models of pregnancy, Interface Focus 9 (5) (2019) 20190026. doi:10.1098/rsfs.2019.0026. URL https://royalsocietypublishing.org/doi/10.1098/rsfs.2019.0026

[26] A. J. Schriefl, T. Schmidt, D. Balzani, G. Sommer, G. A. Holzapfel, Selective enzymatic removal of elastin and collagen from human abdominal aortas: Uniaxial mechanical response and constitutive modeling, Acta Biomaterialia 17 (2015) 125–136. doi:10.1016/j.actbio.2015.01.003. URL https://linkinghub.elsevier.com/retrieve/pii/S1742706115000045

[27] C. E. Barnum, S. S. Shetye, H. Fazelinia, B. A. Garcia, S. Fang, M. Alzamora, H. Li, L. M. Brown, C. Tang, K. Myers, R. Wapner, L. J. Soslowsky, J. Y. Vink, The Non-pregnant and Pregnant Human Cervix: a Systematic Proteomic Analysis, Reproductive Sciences 29 (5) (2022) 1542–1559. doi:10.1007/s43032-022-00892-4. URL https://link.springer.com/10.1007/s43032-022-00892-4

[28] U. o. C. S. D. Allisoon R. Gillies (Department of Bioengineering, University of California San Diego), Richard L. Lieber (Department of Orthopaedic Surgery, Structure and Function of the Skeletal Muscle Extracellular Matrix, Muscle Nerve 44 (3) (2011) 318–331. doi:10.1002/mus.22094.Structure. URL https://www-ncbi-nlm-nih-gov.ezproxy.cul.columbia.edu/pmc/articles/PMC3177172/pdf/nihms277793.pdf

[29] L. Shi, W. Yao, Y. Gan, L. Y. Zhao, W. Eugene McKee, J. Vink, R. J. Wapner, C. P. Hendon, K. Myers, Anisotropic material characterization of human cervix tissue based on indentation and inverse finite element analysis, Journal of Biomechanical Engineering 141 (9). doi:10.1115/1.4043977.

[30] W. Catherino, H. Eltoukhi, A. Al-Hendy, Racial and ethnic differences in the pathogenesis and clinical manifestations of uterine leiomyoma, Seminars in Reproductive Medicine 31 (5) (2013) 370–379. doi:10.1055/s-0033-1348896.

